# Evolution analysis and expression divergence of the chitinase gene family against *Leptosphaeria maculans* and *Sclerotinia sclerotiorum* infection in *Brassica napus*

**DOI:** 10.1101/281923

**Authors:** Wen Xu, Tengsheng Zhou, Bo An, Baojiang Xu, Genyi Li

## Abstract

Blackleg and sclerotinia stem rot caused by *Leptosphaeria maculans* and *Sclerotinia sclerotiorum* respectively are two major diseases in rapeseed worldwide, which cause serious yield losses. Chitinases are pathogenesis-related proteins and play important roles in host resistance to various pathogens and abiotic stress responses. However, a systematic investigation of the chitinase gene family and its expression profile against *L. maculans* and *S. sclerotiorum* infection in rapeseed remains elusive. The recent release of assembled genome sequence of rapeseed allowed us to perform a genome-wide identification of the chitinase gene family. In this study, 68 chitinase genes were identified in *Brassica napus* genome. These genes were divided into five different classes and distributed among 15 chromosomes. Evolutionary analysis indicated that the expansion of the chitinase gene family was mainly attributed to segmental and tandem duplication. Moreover, the expression profiling of the chitinase gene family was investigated using RNA sequencing (RNA-Seq) and the results revealed that some chitinase genes were both induced while the other members exhibit distinct expression in response to *L. maculans* and *S. sclerotiorum* infection. This study presents a comprehensive survey of the chitinase gene family in *B. napus* and provides valuable information for further understanding the functions of the chitinase gene family.

## 1. Introduction

Plant chitinases (EC 3.2.1.14) are enzymes that hydrolyze the N-acetyl glucosamine polymer chitin, a major component of fungal cell walls and exoskeleton of insects (COLLINGE *et al.* 1993) and are considered as one group of pathogenesis-related (PR) proteins (LEGRAND *et al.* 1987), which can be induced in response to the infection of various pathogenic micro-organisms. In the light of classification of glycosyl hydrolases based on amino acid sequence similarities, plant chitinases have been put in glycoside hydrolasis family 18 (GH-18) and 19 (GH-19) (HENRISSAT 1991). According to the CAZy database (http://www.cazy.org/Glycoside-Hydrolases.html) (CANTAREL *et al.* 2009), plant chitinases have been grouped into five different classes ranging from I to V. Of these, classes I, II and IV belong to the GH-19 family whereas the GH-18 family are composed of classes III and V chitinases (HENRISSAT 1991). The details of plant chitinase classification are described as follows. Class I chitinases have an N-terminal chitin-binding domain and a GH-19 catalytic domain. Class II chitinases consist of only a catalytic domain with a high level of sequence and structure similarity to class I chitinases but lack the chitin-binding domain and linker regions. Class IV chitinases show high homology with class I chitinases but are smaller due to one deletion in the chitin-binding domain and three deletions in the catalytic domain (XU *et al.* 2016). Both class III and V chitinases have a GH-18 catalytic domain and a consensus sequence DXDXE, but there is no homology for other amino acids (UMEMOTO *et al.* 2015). GH-18 chitinases are widely distributed in plants, animals, fungi, bacteria and viruses whereas GH-19 members almost exclusively exist in higher plants (PASSARINHO and DE VRIES 2002).

In higher plants, the expression of chitinase genes is involved in defense against biotic and abiotic stress as well as in growth and developmental processes (COLLINGE *et al.* 1993; PUNJA and ZHANG 1993). For instance, a class III chitinase gene, *Mtchitinase III-3*, has been found to be induced upon the infection of fungi *Glomus mosseae* and *Glomus intraradices* in cortical root (BONANOMI *et al.* 2001). PSCHI4, a putative extracellular class II chitinase, is up-regulated in pine seedlings infected with the necrotrophic pathogen *Fusarium subglutinans* f. sp. *Pini* (DAVIS *et al.* 2002). In *Arabidopsis thaliana*, a class IV chitinase gene AtchitIV accumulated very rapidly in leaves after inoculation with *Xanthomonas campestris* and reached maximum mRNA accumulation after one hour infection (GERHARDT *et al.* 1997). In addition, several transgenic studies showed that enhanced levels of chitinase genes in transgenic plants can indeed improve resistance against pathogens and reduce the damage caused by fungi and some insect pests (LIN *et al.* 1995; DING *et al.* 1998; YAMAMOTO *et al.* 2000; WANG *et al.* 2005; PRASAD *et al.* 2013; CHEN *et al.* 2014). There are several reports of induced expression of plant chitinases when plants were exposed to abiotic stresses such as heavy-metal stress (BEKESIOVA *et al.* 2008), drought (HONG and HWANG 2002; LEE *et al.* 2008), salt (HONG and HWANG 2002), cold (YEH *et al.* 2000), heat (KWON *et al.* 2007), UV light and wounding (BREDERODE *et al.* 1991). Furthermore, some chitinases are essential in physiological processes like somatic embryo development (DEJONG *et al.* 1992) and formation of root nodules (OVTSYNA *et al.* 2000). In conclusion, chitinases play important roles in plant defense and plant health.

Rapeseed (*Brassica napus*) is an important oilseed crop worldwide. This crop is affected by various fungal pathogens, especially blackleg caused by *Leptosphaeria maculans* and sclerotinia stem rot by *Sclerotinia sclerotiorum*, which are the most destructive rapeseed diseases in Canada, Australia, Europe and many other regions around the world (WEST *et al.* 2001). Recently, a few studies have been conducted on chitinase genes responding to some pathogens infection in *B. napus* (RASMUSSEN *et al.* 1992a; RASMUSSEN *et al.* 1992b; GRISON *et al.* 1996; WANG *et al.* 2005; AHMED *et al.* 2012). For example, constitutive overexpression of a chimeric chitinase gene in rapeseed had been shown to exhibit an increased resistance to three fungal pathogens compared with their nontransgenic parental plants (GRISON *et al.* 1996). Co-expression of defensin gene *Rs-AFP1* from *R. sativus* and chimeric chitinase gene *chit42* from *T. atroviride* in rapeseed via *Agrobacterium*-mediated transformation demonstrated enhanced resistance against sclerotinia stem rot disease (ZARINPANJEH *et al.* 2016). In addition, global studies of transcriptome dynamics of defense responses to *L. maculans* and *S. sclerotiorum* in *B. napus* presented that pathogen responsive genes including chitinases were rapidly induced during early infection (LOWE *et al.* 2014; HADDADI *et al.* 2016; JOSHI *et al.* 2016; WU *et al.* 2016). However, to date, the chitinase genes in rapeseed have not been systematically identified and thus the genetic resistance to *L. maculans* and *S. sclerotiorum* has been not yet studied. Recently, the availability of the whole genome sequence and RNA-seq sequencing enable further investigations into chitinase genes and their response to *L. maculans* and *S. sclerotiorum* infection on a genome-wide scale (CHALHOUB *et al.* 2014; WOODHOUSE *et al.* 2014).

To further extend the understanding of the chitinase gene family, a global analysis, including identification, sequence features, physical location, the evolutionary relationship and expression pattern of the chitinase gene family in response to *L. maculans* and *S. sclerotiorum* infection in *B. napus* using the RNA-seq sequencing data collected in our lab and some transcriptome data from NCBI database was performed. Expression analysis revealed that some chitinase genes were induced by both pathogens while others displayed differential expression pattern in response to *L. maculans* and *S. sclerotiorum* infection, suggesting that they may have distinct roles in different pathogens stress response. Together, our findings will be helpful for further understanding of the functions of the chitinase gene family against different stress in rapeseed.

## 2. Results

### 2.1 Identification and phylogenetic analysis of the chitinase gene family in *B. napus*

The complete genome sequence and gene annotation was used for the genome-wide identification of the chitinase gene family and a total of 68 putative chitinase genes were identified in the *B. napus* genome (Table 1). All these identified proteins have at least one typical “Glyco_hydro_19” or “Glyco_hydro_18” domain which is responsible for catalyzing the degradation of chitin. Of these, GH-18 family and GH-19 family include 12 and 56 putative chitinase genes, respectively. These chitinase genes in *B. napus* encode proteins ranging from 130 to 1005 amino acids in length with an average of 294. The average number of exons among these chitinase genes was 3.04, a value that is smaller than the average number of exons among all predicted *B. napus* genes (4.9). BLAST search of these 68 proteins against NCBI non-redundant database showed that the top matched hits were endochitinases, chitinases, chitinase-like proteins, which further confirm the reliability of the identified chitinase genes. Furthermore, the signal peptides in 54 predicted chitinase sequences were also identified. To examine the evolutionary relationships among the chitinase genes in *B. napus*, sequence alignment was performed with amino acid sequences (Supplementary Table 1) and an unrooted phylogenetic tree of the 68 chitinase genes using neighbor-joining method was constructed (Figure 1).

**Table 1.** Chitinase genes in the *B.napus* genome and and their sequence characteristics

| Gene | Family | Class | Chr | Start | End | Strand | Protein Length(aa) | Number of Exons | Signal Peptide |
| --- | --- | --- | --- | --- | --- | --- | --- | --- | --- |
| BnaA01g30550D | Glyco_hydro_19 | I | A01 | 20941303 | 20942028 | + | 130 | 1 | YES |
| BnaA01g30560D | Glyco_hydro_19 | I | A01 | 20943578 | 20945861 | + | 145 | 4 | YES |
| BnaA03g32270D | Glyco_hydro_19 | I | A03 | 15569491 | 15570916 | - | 168 | 3 | NO |
| BnaA03g32280D | Glyco_hydro_19 | I | A03 | 15575380 | 15578982 | - | 190 | 6 | YES |
| BnaA05g26640D | Glyco_hydro_19 | I | A05 | 19447511 | 19449216 | + | 196 | 2 | YES |
| BnaC01g38390D | Glyco_hydro_19 | I | C01 | 37477744 | 37480353 | - | 216 | 3 | YES |
| BnaC01g38450D | Glyco_hydro_19 | I | C01 | 37494610 | 37495630 | - | 227 | 2 | NO |
| BnaC03g37570D | Glyco_hydro_19 | I | C03 | 23003071 | 23005099 | - | 231 | 4 | YES |
| BnaC03g37600D | Glyco_hydro_19 | I | C03 | 23016407 | 23018656 | - | 239 | 3 | YES |
| BnaC03g37610D | Glyco_hydro_19 | I | C03 | 23021408 | 23025003 | - | 242 | 2 | YES |
| BnaC05g40680D | Glyco_hydro_19 | I | C05 | 38780461 | 38782151 | + | 245 | 2 | YES |
| BnaA08g31740D | Glyco_hydro_19 | II | A08 | 2107446 | 2110073 | + | 245 | 3 | YES |
| BnaA09g00340D | Glyco_hydro_19 | II | A09 | 146643 | 148245 | - | 245 | 2 | YES |
| BnaA10g01020D | Glyco_hydro_19 | II | A10 | 536566 | 537828 | - | 255 | 2 | YES |
| BnaA10g03880D | Glyco_hydro_19 | II | A10 | 2063411 | 2066519 | - | 255 | 3 | YES |
| BnaC05g01140D | Glyco_hydro_19 | II | C05 | 600283 | 601194 | - | 256 | 2 | YES |
| BnaC05g03990D | Glyco_hydro_19 | II | C05 | 1957654 | 1959846 | - | 257 | 3 | YES |
| BnaC05g36460D | Glyco_hydro_19 | II | C05 | 35739352 | 35741554 | + | 261 | 9 | YES |
| BnaC08g01540D | Glyco_hydro_19 | II | C08 | 1206975 | 1209667 | - | 261 | 3 | YES |
| BnaCnng01370D | Glyco_hydro_19 | II | Cnn | 1527030 | 1528502 | + | 263 | 2 | YES |
| BnaC07g30330D | Glyco_hydro_18 | III | C07 | 34864788 | 34866603 | + | 263 | 3 | YES |
| BnaC09g04600D | Glyco_hydro_18 | III | C09 | 2638283 | 2639657 | + | 263 | 3 | YES |
| BnaA09g05050D | Glyco_hydro_18 | III | A09 | 2477135 | 2478513 | + | 264 | 3 | NO |
| BnaA06g26630D | Glyco_hydro_18 | III | A06 | 18298306 | 18300242 | - | 307 | 3 | YES |
| BnaA03g20290D | Glyco_hydro_19 | IV | A03 | 9648039 | 9649318 | - | 269 | 3 | YES |
| BnaA03g20300D | Glyco_hydro_19 | IV | A03 | 9652001 | 9653298 | - | 269 | 3 | YES |
| BnaA03g20310D | Glyco_hydro_19 | IV | A03 | 9663327 | 9664876 | - | 272 | 2 | YES |
| BnaA03g20320D | Glyco_hydro_19 | IV | A03 | 9678956 | 9680269 | - | 272 | 2 | YES |
| BnaA03g20330D | Glyco_hydro_19 | IV | A03 | 9684457 | 9685621 | - | 274 | 2 | YES |
| BnaA03g20340D | Glyco_hydro_19 | IV | A03 | 9698939 | 9700499 | - | 275 | 2 | YES |
| BnaA03g56430D | Glyco_hydro_19 | IV | A03 | 711494 | 712848 | + | 275 | 2 | YES |
| BnaA04g25220D | Glyco_hydro_19 | IV | A04 | 18233599 | 18234851 | - | 275 | 2 | NO |
| BnaA04g25230D | Glyco_hydro_19 | IV | A04 | 18235520 | 18236868 | - | 279 | 2 | NO |
| BnaA05g03420D | Glyco_hydro_19 | IV | A05 | 1888936 | 1890632 | - | 279 | 3 | YES |
| BnaA05g03430D | Glyco_hydro_19 | IV | A05 | 1893397 | 1894609 | - | 280 | 2 | YES |
| BnaA05g03440D | Glyco_hydro_19 | IV | A05 | 1904615 | 1905892 | - | 280 | 2 | YES |
| BnaA05g08640D | Glyco_hydro_19 | IV | A05 | 4802795 | 4804226 | + | 281 | 2 | YES |
| BnaA09g15430D | Glyco_hydro_19 | IV | A09 | 8977793 | 8978880 | + | 281 | 2 | YES |
| BnaA09g15440D | Glyco_hydro_19 | IV | A09 | 8979421 | 8980691 | + | 281 | 2 | NO |
| BnaA09g34290D | Glyco_hydro_19 | IV | A09 | 25164422 | 25167344 | + | 281 | 6 | YES |
| BnaA10g05680D | Glyco_hydro_19 | IV | A10 | 3646850 | 3650151 | - | 282 | 2 | YES |
| BnaC03g19370D | Glyco_hydro_19 | IV | C03 | 10063312 | 10065182 | - | 282 | 2 | YES |
| BnaC03g24270D | Glyco_hydro_19 | IV | C03 | 13607893 | 13609216 | - | 282 | 2 | YES |
| BnaC03g24280D | Glyco_hydro_19 | IV | C03 | 13614270 | 13615575 | - | 282 | 3 | YES |
| BnaC03g24290D | Glyco_hydro_19 | IV | C03 | 13628566 | 13630189 | - | 283 | 2 | YES |
| BnaC03g24300D | Glyco_hydro_19 | IV | C03 | 13641790 | 13643090 | - | 284 | 2 | YES |
| BnaC03g24310D | Glyco_hydro_19 | IV | C03 | 13644193 | 13644910 | - | 302 | 2 | YES |
| BnaC03g24330D | Glyco_hydro_19 | IV | C03 | 13646581 | 13649085 | - | 302 | 2 | YES |
| BnaC03g24340D | Glyco_hydro_19 | IV | C03 | 13654943 | 13656106 | - | 302 | 2 | NO |
| BnaC03g24360D | Glyco_hydro_19 | IV | C03 | 13695822 | 13697335 | - | 318 | 2 | YES |
| BnaC04g09720D | Glyco_hydro_19 | IV | C04 | 7384173 | 7385277 | + | 319 | 5 | YES |
| BnaC04g48920D | Glyco_hydro_19 | IV | C04 | 47325431 | 47326573 | + | 320 | 2 | YES |
| BnaC04g49090D | Glyco_hydro_19 | IV | C04 | 47408585 | 47409809 | - | 322 | 2 | NO |
| BnaC04g49100D | Glyco_hydro_19 | IV | C04 | 47411832 | 47413177 | - | 322 | 2 | YES |
| BnaC04g53030D | Glyco_hydro_19 | IV | C04 | 680771 | 682319 | - | 322 | 3 | YES |
| BnaC04g53040D | Glyco_hydro_19 | IV | C04 | 688006 | 689125 | - | 322 | 2 | NO |
| BnaC08g25190D | Glyco_hydro_19 | IV | C08 | 27018000 | 27020278 | + | 323 | 6 | NO |
| BnaC08g25210D | Glyco_hydro_19 | IV | C08 | 27024727 | 27025633 | + | 329 | 2 | NO |
| BnaC09g51720D | Glyco_hydro_19 | IV | C09 | 948730 | 950079 | - | 334 | 3 | YES |
| BnaCnng39650D | Glyco_hydro_19 | IV | Cnn | 38235388 | 38236682 | + | 342 | 3 | YES |
| BnaC03g63440D | Glyco_hydro_18 | V | C03 | 52860028 | 52861720 | + | 346 | 3 | NO |
| BnaA01g34980D | Glyco_hydro_18 | V | A01 | 304675 | 306686 | + | 363 | 3 | YES |
| BnaUnng03570D | Glyco_hydro_18 | V | Unn | 5067935 | 5068651 | - | 371 | 1 | NO |
| BnaA03g26800D | Glyco_hydro_18 | V | A03 | 13187584 | 13189629 | + | 381 | 8 | YES |
| BnaA03g26330D | Glyco_hydro_18 | V | A03 | 12888937 | 12890809 | + | 383 | 3 | YES |
| BnaC03g31740D | Glyco_hydro_18 | V | C03 | 19525415 | 19530521 | + | 424 | 18 | YES |
| BnaA08g09330D | Glyco_hydro_18 | V | A08 | 8944530 | 8946552 | - | 513 | 3 | YES |
| BnaC01g41020D | Glyco_hydro_18 | V | C01 | 240723 | 242688 | + | 1005 | 5 | YES |

**Figure 1.**
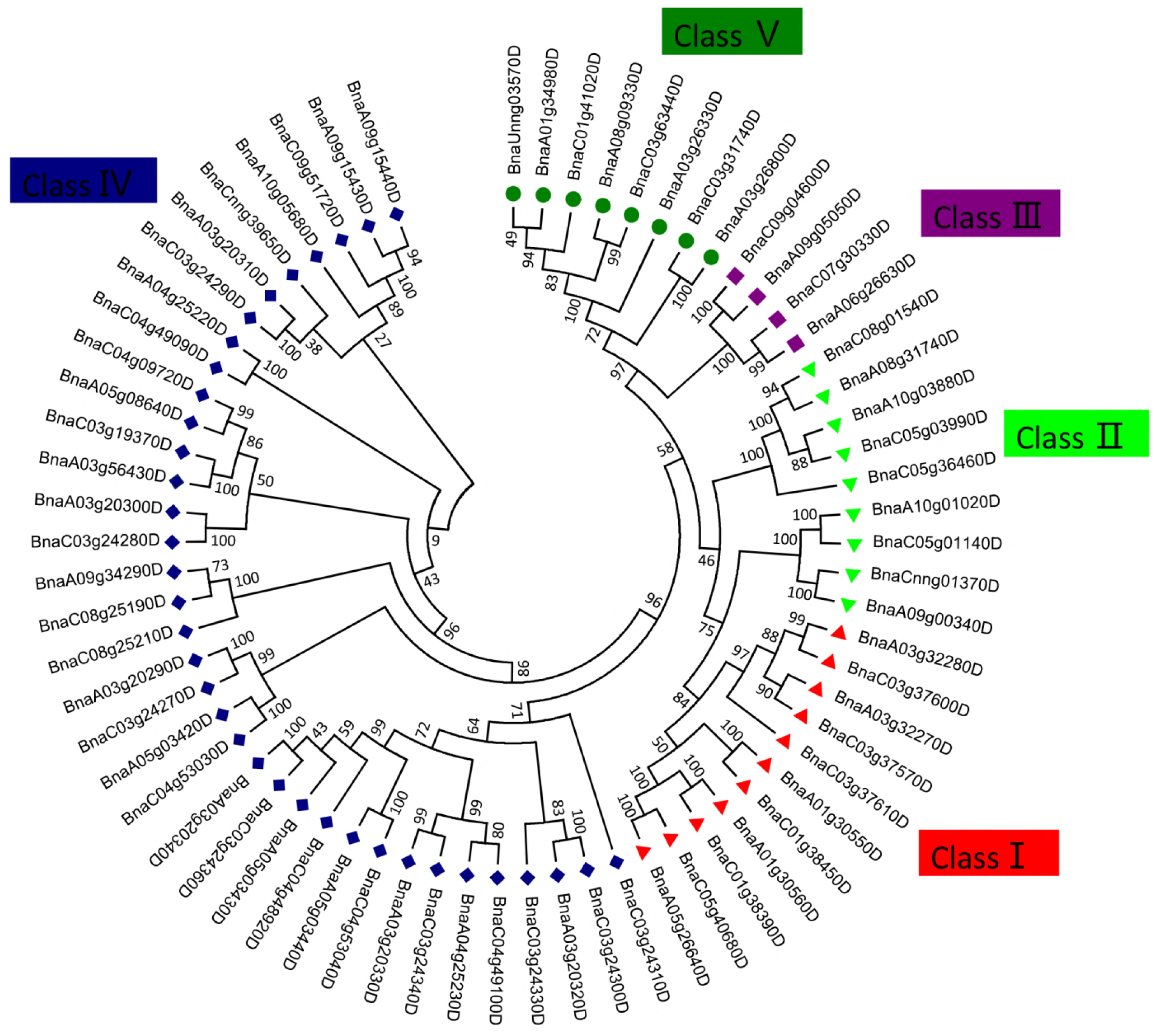
Phylogenetic tree of the chitinase gene family in *Brassica napus*. The chitinase protein sequences were used to construct multiple sequence alignments using MUSCLE program within MEGA 7.0 software. Phylogenetic analysis was performed using MEGA 7.0 with the neighbor-joining method with 1,000 bootstrap replications.

### 2.2 Conserved motifs and gene structures of the chitinase gene family in *B. napus*

The above phylogenetic tree highlighted that the 68 chitinase genes could be divided into five well-supported subfamilies, which were consistent with Class I, II, III, IV, and V. As expected, the chitinase genes of Glyco_hydro_19 and Glyco_hydro_18 families were clustered into two relatively distinct branches. Chitinase genes from subfamilies Class III and Class V are in the Glyco_hydro_18 clade, whereas subfamilies Class I, II, and IV that belong to Glyco_hydro_19 clade were clustered together and showed close relationships (Figure 2). According to the phylogenetic tree, chitinases in different subfamilies had various characteristics. Among the five subfamilies, Class IV was found to be the biggest group with 36 members, accounting for almost a half of the chitinase gene family, whereas there were 11, 9, 4, 8 chitinase genes in Class I, II, III, V subfamilies, respectively.

**Figure 2.**
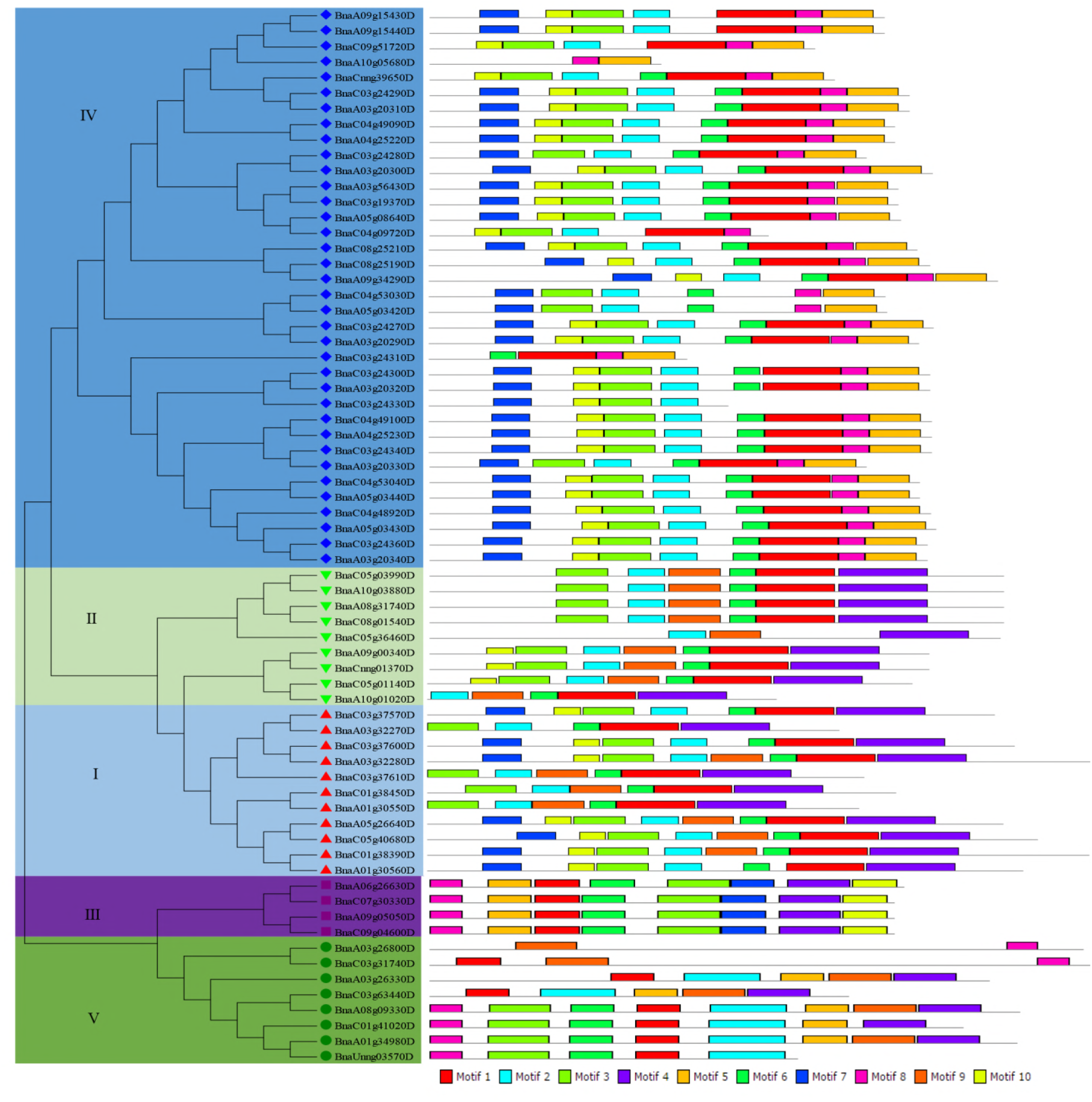
Phylogenetic relationships and motif compositions of chitinase genes. The phylogenetic tree was created in MEGA7.0 software. Five major phylogenetic groups designated as I to V were marked with different color backgrounds. Schematic representation of the conserved motifs of the chitinase gene family was elucidated by MEME software. Each motif was represented by a colored box numbered in the bottom. The details of individual motif were shown in Supplementary Figure 1 and 2.

To gain further insights into the structural diversity and functional evolution of chitinase genes, 10 motifs of all 68 chitinase genes are captured by MEME software and displayed schematically in Figure 2 and Supplementary Figure 1and 2. As shown in Figure 2, most chitinases in the same class shared common motif compositions. Among the GH-19 family, motifs 1,3,4,9 were annotated as glycoside hydrolase catalytic domain and motif 7 was annotated as chitin-binding domain. Each member of chitinases in classes I, II and IV had at least one glycoside hydrolase catalytic domain and motif 4 was only detected in classes I and II. Most chitinase genes from subfamilies I and IV also harbored a chitin-binding domain whereas motif 7 did not exist in the subfamily II. In the GH-18 family, except for motif 8 and 9, other 8 motifs were annotated as glycoside hydrolase catalytic domains. Similarly, in the GH-19, at least one glycoside hydrolase catalytic domain was detected in every chitinase gene of class III and V. Interestingly, motif 7 and motif 10 are unique in class III whereas motif 2 can be only detected in class V. In addition, we analyzed the coding sequences with corresponding genome sequences of each chitinase genes in *B. napus*. A detailed illustration of the chitinase gene structure was shown in Figure 3. Most chitinase genes contained two or three exons, whereas three chitinase genes (BnaA01g30550D, BnaA0626330D and BnaUnng03570D) had no introns. In general, most chitinase genes in the same class showed similar conserved motifs and exon-intron structures. These findings revealed that motif compositions and gene structures of each class in the chitinase gene family were relatively conserved. The similar features of chitinase genes in the same class may fulfill similar functions and this claim need to be supported by their expression and related data.

**Figure 3.**
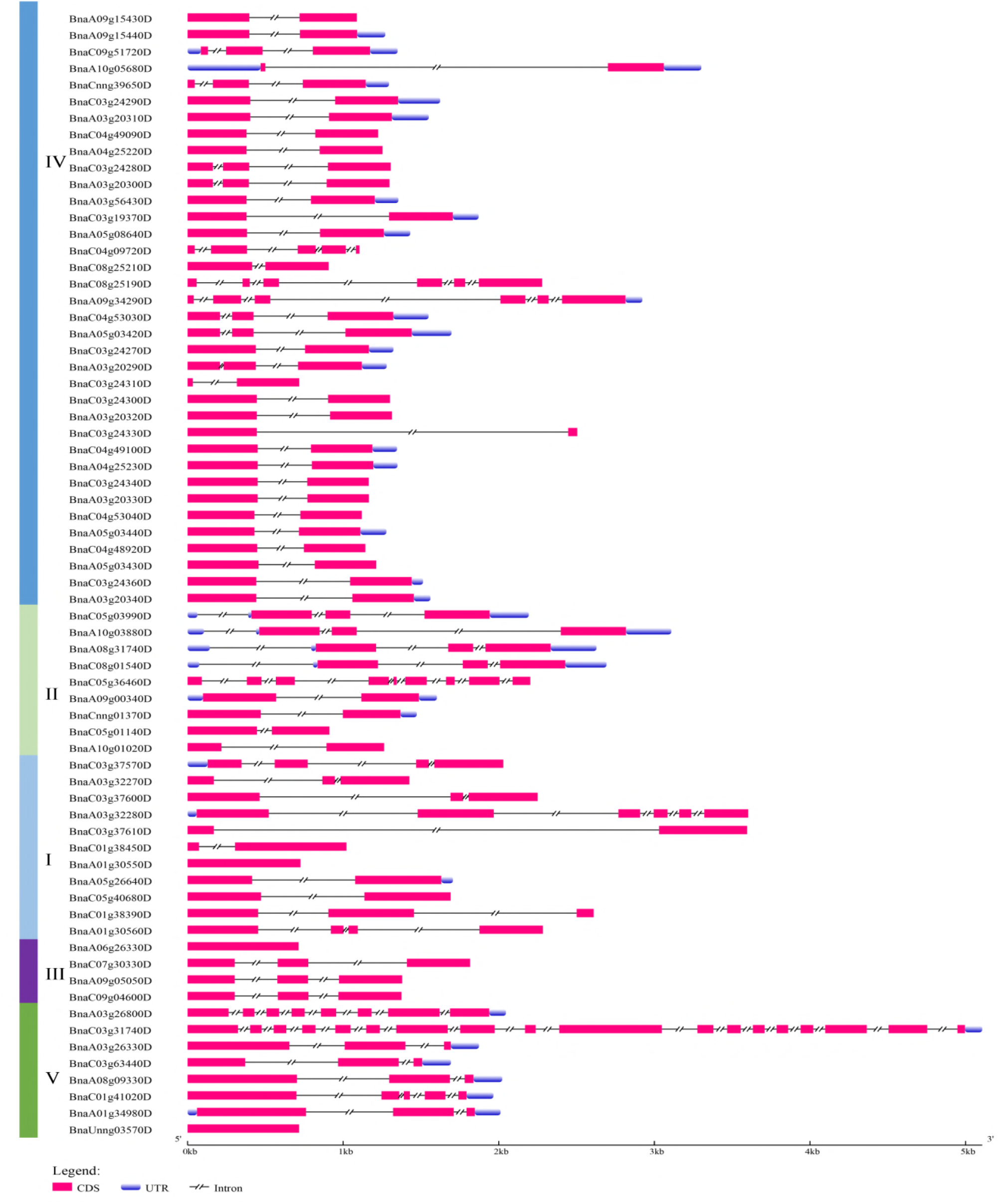
Exon-intron structures of all chitinase genes in *Brassica napus*. Schematic diagram represents the gene structure of all 68 chitinase genes identified in this study using Gene Structure Display Server (http://gsds.cbi.pku.edu.cn/). CDS are shown as red boxes; introns are indicated by double slashes on the bar; UTR sequences are shown as blue boxes.

### 2.3 Chromosomal distribution and evolution patterns of chitinase genes in *B. napus*

The chromosomal distribution of 68 chitinase genes was analyzed based on the available gene annotation and genome sequence assembly. The results revealed that all 68 chitinase genes were distributed among 15 out of 19 chromosomes with the exception of chromosomes A02, A07, C02 and C06 in the *B. napus* genome (Figure 4a). There were 32 genes mapped in the A genome, and 35 genes located in the C genome while one gene BnaUnng03570D was not assigned to a chromosome. GH-19 family presented on 13 chromosomes except chromosomes A06 and C07 while GH-18 family were absent from chromosomes A04, A05, A10, C04, C05, C08. The number of chitinase genes varied considerably among different chromosomes and the large numbers of chitinase genes were found on chromosomes A03 and C03, harbouring 11 and 14 genes, respectively (Figure 4a). Furthermore, the classes of the chitinase gene family were distributed in different chromosomes and up to four classes in A03 and C03 chromosomes were identified (Figure 4b)

**Figure 4.**
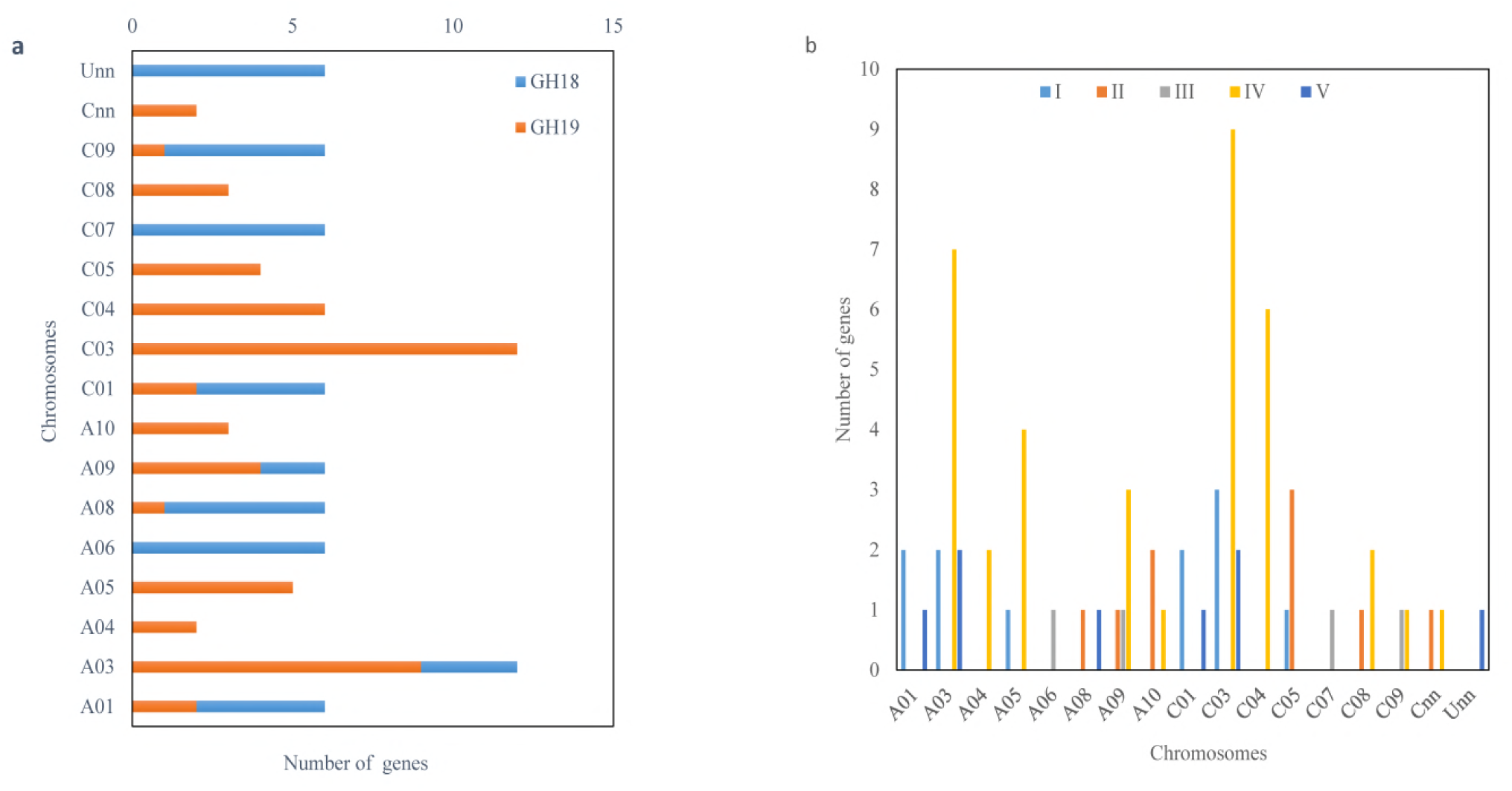
Distribution of the chitinase gene family on *Brassica napus* chromosomes. (a) Gene distribution of GH-18 family and GH-19 family on *Brassica napus* chromosomes. (b) Distribution of five subfamilies of chitinase genes on *Brassica napus* chromosomes.

To understand the evolution of the chitinase gene family, twenty-six pairs of paralogs were detected in 68 chitinase genes based on criteria for both coverage ≥70% and identity ≥70% (Table 2). The phylogenetic relationship analysis of chitinase genes also showed that most pairs of paralogs could be clustered together (Figure 1). For example, the four members in two pairs of paralogs (BnaA09g05050D and BnaC09g04600D, BnaA06g26630D and BnaC07g30330D) in Class III subfamily were clustered into two parts in a single clade. As genome duplication was considered, one member in the A subgenome would correspond to one homologous gene in the C subgenome in *B. napus*. In fact, 50 members of 68 chitinase genes showed such a one-to-one correspondence.

**Table 2.** Table listing the chitinase genesparalog sets among A and C subgenomes of *B. napus* and their orthologs in *A. thaliana, B.rapa, B.oleracea*.

| *B.napus_C* vs<br>*A. thaliana* | *B.napus_A* vs *A.*<br>*thaliana* | *B.napus_C* vs<br>*B.oleracea* | *B.napus_A* vs<br>*B.rapa* | *B.napus_A* | *B.napus_C* |
| --- | --- | --- | --- | --- | --- |
|  |  |  |  | BnaA03g32270D | BnaC03g37570D |
| AT2G43610.1 | AT2G43610.1 | Bo3g036710.1 | Bra000314.1 | BnaA03g20330D | BnaC03g24340D |
| AT2G43620.1 | AT2G43620.1 | Bo3g036730.1 | Bra000315.1 | BnaA03g20340D | BnaC03g24360D |
| AT2G43590.1 | AT2G43590.1 | Bo3g036660.1 | Bra000311.1 | BnaA03g20310D | BnaC03g24290D |
|  |  |  | Bra027940.1 | BnaA09g15430D | BnaA09g15440D |
|  |  |  | Bra027940.1 | BnaA09g15440D | BnaA09g15430D |
| AT4G01700.1 | AT4G01700.1 | Bo9g004220.1 | Bra036316.1 | BnaA09g00340D | BnaCnng01370D |
| AT4G19810.1 | AT4G19810.1 | Bo1g021980.1 | Bra020951.1 | BnaA08g09330D | BnaC03g63440D |
| AT5G24090.1 | AT5G24090.1 | Bo9g013820.1 | Bra026469.1 | BnaA09g05050D | BnaC09g04600D |
|  |  | Bo3g036670.1 | Bra000312.1 | BnaA03g20320D | BnaC03g24300D |
|  |  | Bo3g028270.1 | Bra023009.1 | BnaA03g56430D | BnaC03g19370D |
| AT3G12500.1 | AT3G12500.1 | Bo5g133420.1 | Bra034754.1 | BnaA05g26640D | BnaC05g40680D |
| AT2G43580.1 | AT2G43580.1 | Bo3g036650.1 | Bra000311.1 | BnaA03g20300D | BnaC03g24280D |
|  | AT2G43590.1 | Bo4g194450.1 | Bra037699.1 | BnaA04g25220D | BnaC04g49090D |
|  |  | Bo5g003080.1 | Bra033318.1 | BnaA10g01020D | BnaC05g01140D |
|  |  | Bo4g020930.1 | Bra004773.1 | BnaA05g03440D | BnaC04g53040D |
|  |  | Bo1g139190.1 | Bra038727.1 | BnaA01g30550D | BnaC01g38450D |
| AT2G43610.1 | AT2G43610.1 | Bo4g194460.1 | Bra037698.1 | BnaA04g25230D | BnaC04g49100D |
| AT2G43570.1 | AT2G43570.1 | Bo3g036640.1 | Bra000310.1 | BnaA03g20290D | BnaC03g24270D |
| AT5G24090.1 | AT5G24090.1 | Bo7g097210.1 | Bra009731.1 | BnaA06g26630D | BnaC07g30330D |
| AT1G05850.1 | AT1G05850.1 | Bo8g005830.1 | Bra030628.1 | BnaA08g31740D | BnaC08g01540D |
| AT4G19810.1 | AT4G19810.1 | Bo1g021960.1 | Bra013426.1 | BnaA01g34980D | BnaC01g41020D |
|  |  | Bo8g083650.1 |  | BnaA09g34290D | BnaC08g25190D |
| AT1G05850.1 | AT1G05850.1 | Bo5g007040.1 | Bra015457.1 | BnaA10g03880D | BnaC05g03990D |
|  | AT3G12500.1 | Bo1g139210.1 | Bra038726.1 | BnaA01g30560D | BnaC01g38390D |
| AT2G43570.1 |  | Bo4g020920.1 | Bra004771.1 | BnaA05g03420D | BnaC04g53030D |
| AT2G43620.1 | AT2G43620.1 | Bo3g036730.1 | Bra004772.1 | BnaA05g03430D | BnaC04g48920D |
|  | AT4G19750.1 |  | Bra000892.1 | BnaA03g26330D |  |
|  | AT4G01040.1 |  | Bra000939.1 | BnaA03g26800D |  |
|  |  |  | Bra001454.1 | BnaA03g32280D |  |
|  | AT2G43590.1 |  | Bra005345.1 | BnaA05g08640D |  |
|  |  |  | Bra039597.1 | BnaA10g05680D |  |
| AT2G43590.1 |  | Bo3g024590.1 |  |  | BnaCnng39650D |
|  |  | Bo9g055720.1 |  |  | BnaC09g51720D |
| AT3G54420.1 |  | Bo8g083670.1 |  |  | BnaC08g25210D |
|  |  | Bo5g123590.1 |  |  | BnaC05g36460D |
|  |  |  |  |  | BnaC04g09720D |
|  |  | Bo3g064970.1 |  |  | BnaC03g37610D |
|  |  |  |  |  | BnaC03g37600D |
|  |  | Bo3g054420.1 |  |  | BnaC03g31740D |
|  |  | Bo3g036700.1 |  |  | BnaC03g24330D |
| AT2G43600.1 |  | Bo3g036680.1 |  |  | BnaC03g24310D |
| AT4G19810.1 |  | Bo1g021980.1 | Bra020951.1 |  | BnaUnng03570D |

The distribution of chitinase genes indicated a relatively deep evolutionary origin of these chitinase genes as well as gene duplication. Previous research suggests that the evolution of a plant disease resistance gene family is usually mediated by recombination, tandem duplication, and segmental duplication (LEISTER 2004). The allotetraploid *B. napus* is a spontaneous hybridisation of *B. rapa* (A genome) and *B. olearecea* (C genome). To understand the origin and duplication patterns of these chitinase genes, putative orthologs of chitinase genes in *B. napus* were also identified in *A. thaliana, B. rapa* and *B. olearecea* (Table 2). The results demonstrated that 30 orthologs of 32 chitinase genes in the A subgenome were identified in *B. rapa* and 32 orthologs of 35 chitinase genes in the C subgenome in *B. olearecea.* The gene BnaUnng03570D had both orthologs (Bo1g021980.1 and Bra020951.1) in *B. rapa* and *B. olearecea*, respectively, but it had much higher identities with Bo1g021980.1. Furthermore, the order and synteny of chitinase genes BnaA03g20290D, BnaA03g20300D, BnaA03g20310D, BnaA03g20320D, BnaA03g20330D, BnaA03g20340D in chromosome A03 and BnaC03g24270D, BnaC03g24280D, BnaC03g24290D, BnaC03g24300D, BnaC03g24340D, BnaC03g24360D in chromosome C03, as well as those genes for both subgenomes of the allopolyploid (the A and C subgenomes in *B. napus* vs. the A and C genomes in *B. rapa* and *B. olearecea*) revealed that there were a few structural rearrangements and genomic collinearity in *B. napus* with regard to the chitinase gene family evolution and expansion. These findings showed that most chitinase genes (63/68) of allotetraploid *B. napus* were inherited from their diploid ancestors by recombination or segmental duplication. It is interesting to observe that the best hit of chitinase gene BnaA09g15430D was BnaA09g15440D which were tandem repeats on the A09 chromosome. It was also observed that five chitinase genes in *B. napus* had no orthologs in their diploid ancestors. Of these, three genes (BnaA09g34290D, BnaC04g09720D and BnaC03g37600D) and one paralog of two genes (BnaA03g32270D and BnaC03g37570D) did not have orthologs in *A. thaliana, B. rapa* and *B. olearecea*, which might result from incomplete and error-filled genome assemblies and gene annotation errors or gene structure rearrangement in the evolutionary process. Perhaps these five genes might be the new members of the chitinase gene family during the evolution in *B. napus*.

### 2.4 Transcriptomic profiles of the chitinase gene family in response to *L. maculans* and *S. sclerotiorum* during early infection in *B. napus*

Blackleg, also known as stem canker caused by *L. maculans* and sclerotinia stem rot caused by *S. sclerotiorum* are two major rapeseed diseases in most major rapeseed growing areas. The expression profiling of all chitinase genes in response to *L. maculans* and *S. sclerotiorum* infection in resistant and susceptible *B. napus* accessions were investigated using whole-transcriptome sequencing data to understand the roles of chitinase genes at early stages of infection. The results showed that these two biostresses caused a significant expression induction of some members in the chitinase gene family. The differentially expressed genes (DEGs) of chitinases in response to *L. maculans* were identified. Among all 68 chitinase genes in the genome, 31 and 25 chitinase genes were up-regulated in the resistant lines and susceptible lines inoculated with the pathogen compared with their water control, respectively. Of these up-regulated chitinase genes, 24 were overlapped in the resistant lines and susceptible accessions. Combined data of all resistant and susceptible lines revealed that 15 chitinase genes were upregulated and no chitinase gene was dramatically downregulated during the infection of pathogen *L. maculans* (Figure 5).

**Figure 5.**
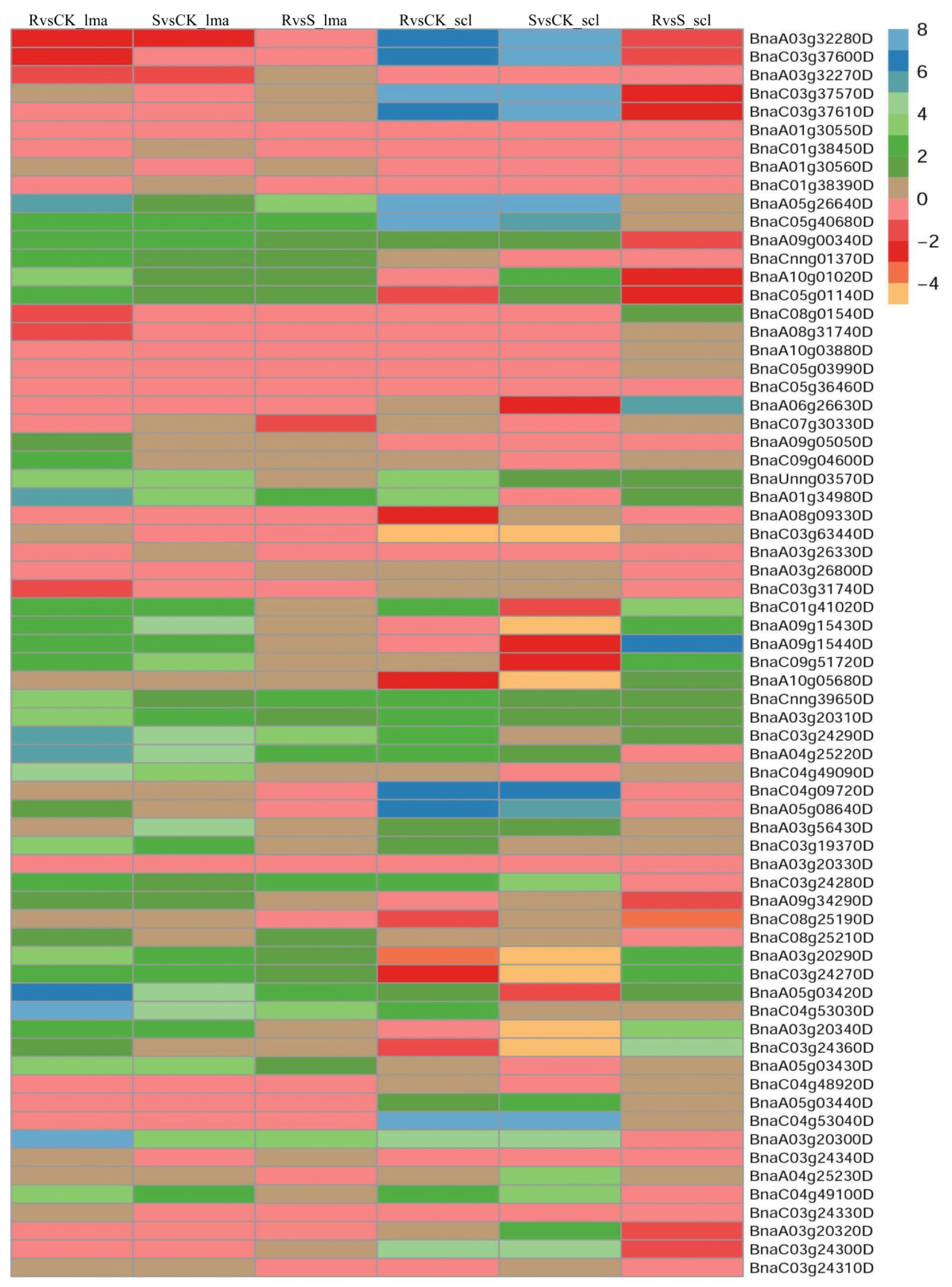
Expression profiles of chitinase genes in response to *Leptosphaeria maculans* (lma) and *Sclerotinia sclerotiorum* (scl) infection. Relative fold change in as compared to controlin in resistant and susceptible *B. napus* lines and was used to generate heatmap. R and S were represented as resistant and susceptible lines, respectively. The colored scale for the relative expression levels is shown.

In a previous report, the abundance of transcripts in resistant and susceptible *B. napus* accessions at the 4 day post-inoculation treatments were analyzed to understand the differential defense response to *S. sclerotiorum* (WU *et al.* 2016). Using these data and all 68 chitinase genes identified in this study, 23 and 20 chitinase genes were up-regulated while 6 and 11 members were downregulated in the resistant accession and susceptible accession compared with water control, respectively. Compared with the expression of chitinase genes in response to blackleg pathogen infection, 16 up-regulated and 4 down-regulated chitinase genes were overlapped respectively. Compared with the S accession, the analyses showed that 13 and 7 chitinase genes were the same as those upregulated and downregulated ones against *S. sclerotiorum* attack, respectively.

Chitinase genes which were induced after infection with *L. maculans* and *S. sclerotiorum* in rapeseed were further classified into three groups (Table 3). In the first group, 10 members of the chitinase gene family had stronger up-regulation in resistant accessions than susceptible counterparts with more than two values of log2. These 10 chitinase genes showed a wide range of upregulation in resistant accession infected by *S. sclerotiorum* and the differences of the upregulation between resistant and susceptible accessions varied and were less than two in the log2 values. There were 9 chitinase genes showing higher upregulation in the infection of resistant accessions with *L. maculans* with the log2 values of more that three while most of these genes were less upregulated in susceptible accessions. In the infection of *S. sclerotiorum*, only few chitinase genes in the second group showed upregulated. Eight members of chitinase genes in the third group showed very high levels of upregulation with the log2 values ranging from 4.45-7.96 in the infection of *S. sclerotiorum* while the differences in both resistant and susceptible accessions were little or even higher in susceptible accessions. The chitinase genes in the third group showed much less variation in the infection of *L. maculans* and the expression of some of these genes were not detected, indicating that these genes did not play a critical role in the defense against *L. maculans.* The results suggested that the cross-talk under different pathogen attack and redundant functions can be maintained over long evolutionary periods. On the other hand, eight chitinase genes were preferentially induced by *L. maculans* infection while other 8 chitinase genes were preferentially increased by *S. sclerotiorum* infection respectively. The distinctive expression pattern of the chitinase gene family suggested that the functions of different members in the chitinase gene family have diverged during long-term evolution and might exhibit different roles against different biotic and abiotic stresses.

**Table 3.**
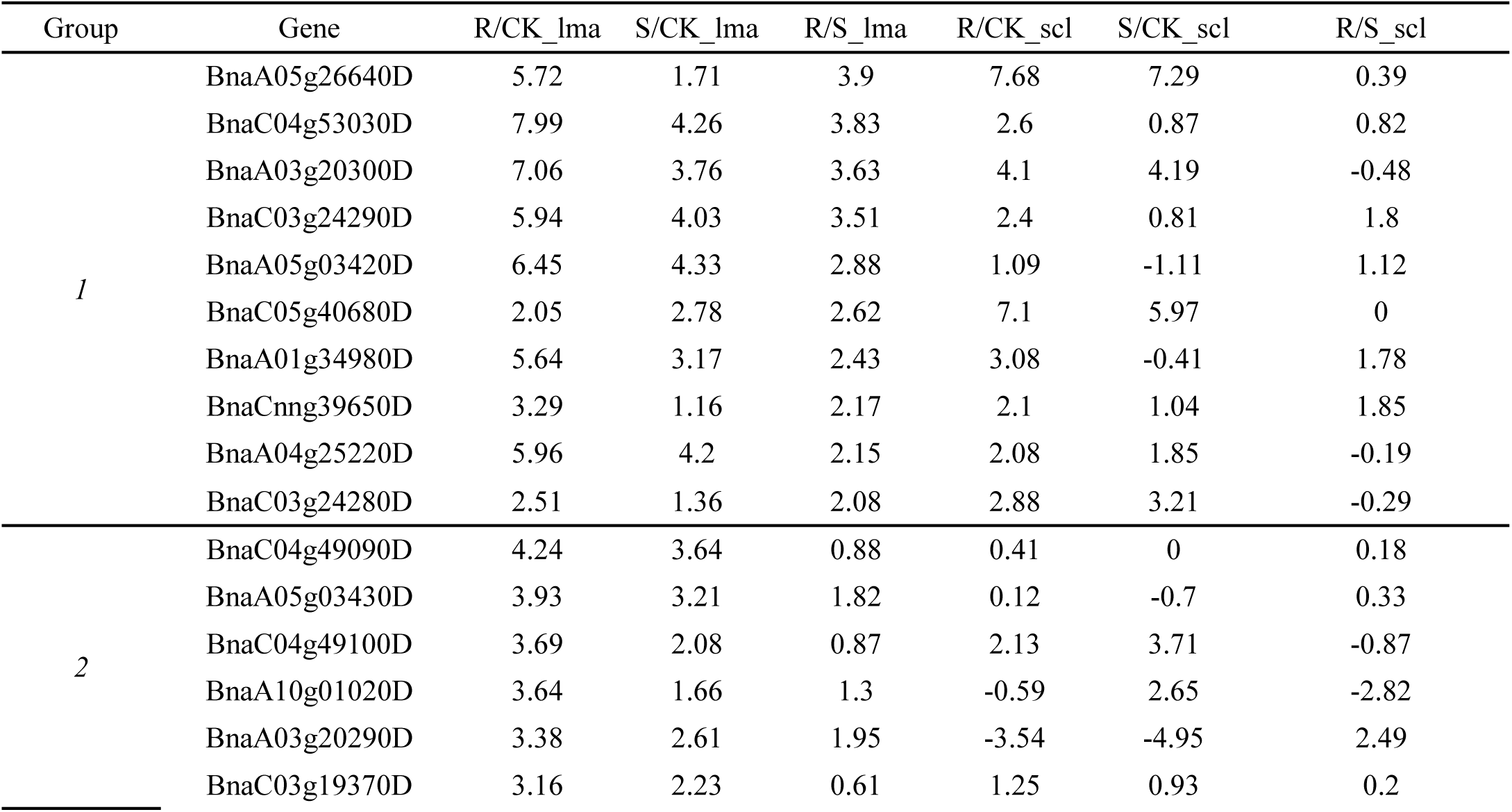
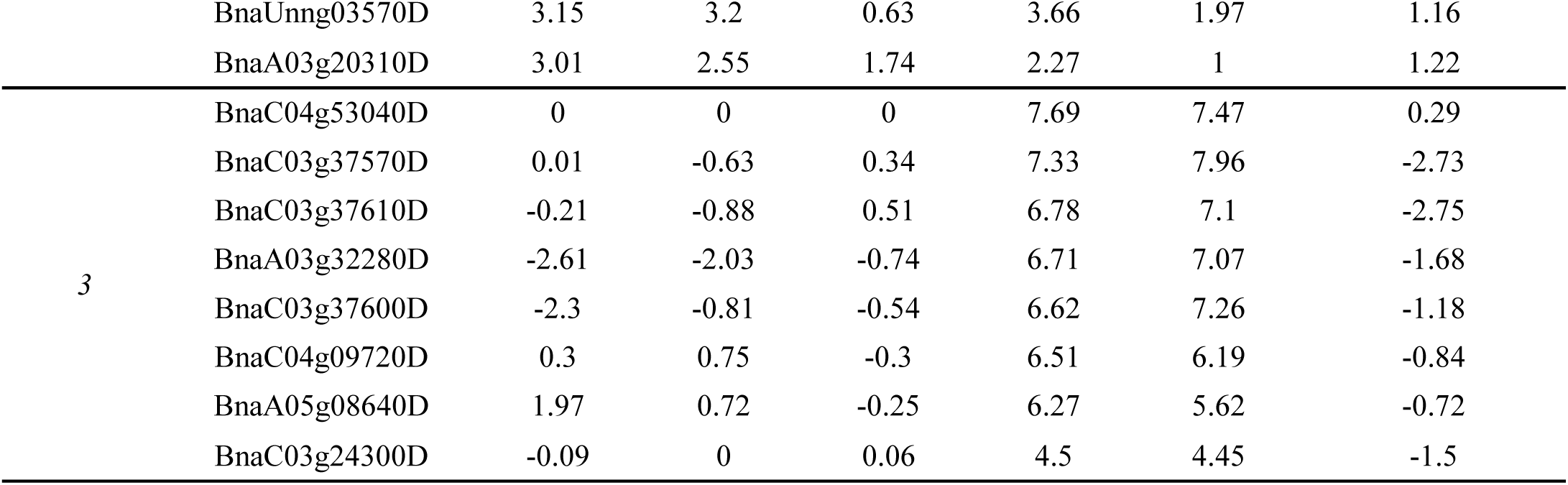
Fold change (log2) of chitinase gene expression in response to *Leptosphaeria maculans* (lma) and *Sclerotinia sclerotiorum* (scl) infection in resistant and susceptible *B. napus* lines.

## 3. Discussion

Plants have developed highly sophisticated immune mechanisms to respond to pathogen attack by the induction of expression of a large number of genes encoding pathogenesis-related (PR) proteins, such as chitinases. Chitinases are believed to play important roles in plant-pathogen interactions and catalyze the hydrolysis of the β-1-4-linkage in the N-acetyl-D-glucosamine polymer of chitin, which is a major component of many fungal cell walls, but absent in higher plants (LEGRAND *et al.* 1987; COLLINGE *et al.* 1993). The chitinase gene family has been widely characterized as excellent candidates to improve plants tolerance to stresses, including drought (HONG and HWANG 2002; LEE *et al.* 2008), salt (HONG and HWANG 2002), cold (YEH *et al.* 2000), heat (KWON *et al.* 2007), UV light, wounding (BREDERODE *et al.* 1991), fungal pathogens and some insect pests (LIN *et al.* 1995; DING *et al.* 1998; YAMAMOTO *et al.* 2000; WANG *et al.* 2005; PRASAD *et al.* 2013; CHEN *et al.* 2014). However, the genome-wide identification and expression pattern of the chitinase gene family in response to *L. maculans* and *S. sclerotiorum* infection has not been reported in *B. napus*.

In this study, a total of 68 chitinase genes were identified in *B. napus* genome. Of these, GH-18 family and GH-19 family have 12 and 56 chitinase genes, respectively, which was further supported by analysis of gene structure and conserved motifs (Figure 2, 3). GH-18 family was divided into Class III (4 genes) and Class V (8 genes). GH-19 family was composed of Class I (11 genes), Class II (9 genes) and Class IV (36 genes). However, there were 13 and 26 chitinase genes in GH-19 family and GH-18 family in para rubber tree, respectively (MISRA 2015). Class IV had the most members in *B. napus* whereas Class III posed the most genes of chitinase in Para rubber tree, which may reveal that there is evolutionary divergence of specific classes of chitinases in different species. The chitinase gene family has 24, 32, 35, and 68 members in *A. thaliana, B. rapa, B. oleracea*, and *B. napus*, respectively (XU *et al.* 2007), which suggested that chitinase genes in *B. napus* had expanded in comparison to its ancestors. Gene duplication events, such as tandem duplication and segmental duplication play important roles in the rapid expansion and evolution of gene families (XU *et al.* 2012). The AACC genome of *B. napus* was formed through recent allopolyploidy between the ancestors of *B. rapa* (AA) and *B. oleracea* (CC) (CHALHOUB *et al.* 2014). From our analysis, 30 orthologs of 32 chitinase genes in the A genome were identified in *B. rapa* and 32 orthologs of 35 chitinase genes in the C genome of *B. olearecea*. Most orthologous gene pairs in *B. rapa* and *B. oleracea* are still homeologous pairs in *B. napus*. Most chitinase genes in *B. napus* showed a close relationship to their ancestor chitinase genes, which suggested that segmental duplication or polyploidy events contributed to the expansion of the chitinase gene family in *B. napus*. Only one pair of tandemly duplicated genes (BnaA09g15430D and BnaA09g15440D) was identified. These findings suggest that segmental duplication and tandem duplication likely plays an important role in the expansion of the chitinase gene family in *B. napus*. Moreover, five genes (BnaA09g34290D, BnaC04g09720D, BnaC03g37600D, BnaA03g32270D and BnaC03g37570D) do not have orthologs in *A. thaliana, B. rapa* and *B. olearecea*, suggesting that they may be the new members of the chitinase gene family and coevolve with fungi in response to variation in pathogen defenses.

Chitinases play a major role in host defense by directly attacking fungal pathogens in *A. thaliana* (GERHARDT *et al.* 1997), rice (LIN *et al.* 1995), grapevine (YAMAMOTO *et al.* 2000), tobacco (DING *et al.* 1998; CHEN *et al.* 2014; DONG *et al.* 2017), peanut (PRASAD *et al.* 2013) and pepper (HONG and HWANG 2002). Some chitinase members can be induced by fungal pathogens, such as *Cylindrosporium concentricum, Phoma lingam*, and *S. sclerotiorum*. The role of chitinase gene also had been studied in *B. napus* and overexpression of chitinase genes could increase tolerance in transgenic plants previously (GRISON *et al.* 1996; ZARINPANJEH *et al.* 2016). Transgenic plants of *B. napus* cv. ZS 758 carrying sporamin and chitinase PjChi-1 genes exhibited increased levels of resistance to *S. sclerotiorum* and reduced the size of leaf spot in transformants compared to untransformed wild-type plants (LIU *et al.* 2011). However, constitutive expression of pea chitinase gene showed little or no enhancement of resistance to *L. maculans* in transgenic rapeseed compared with non-expressing transgenic lines (WANG *et al.* 1999). Some chitinases such as pineapple leaf chitinase-A do not have any antifungal activity (TAIRA *et al.* 2005). Although many studies about the individual member of the chitinase gene family have been published, there is little information about analysis of their expression divergence of the chitinase gene family at a genome-wide level in rapeseed, especially under different fugal pathogen stresses. In this study, detailed expression pattern of the chitinase gene family against *L. maculans* and *S. sclerotiorum* infection in rapeseed was analyzed using RNA-seq data. The results showed that many chitinase genes could transcriptionally respond to *L. maculans* and *S. sclerotiorum* infection in rapeseed (Figure 5), implying possible function of chitinase genes in response to these two fugal pathogens in *B. napus*. The results reveal that the resistant accesions differentiate from the susceptible ones in pathogen defense so we hypothesized that the function of different chitinase genes has been diverged against pathogens in resistant and susceptible *B. napus* accessions. Previously, the expression of chitinase genes could be induced in response to all kinds of pathogens, such as *G. mosseae, F. subglutinans* f. sp. *Pini, X. campestris,L. maculans* and *S. sclerotiorum* (GERHARDT *et al.* 1997; BONANOMI *et al.* 2001; DAVIS *et al.* 2002; LOWE *et al.* 2014; WU *et al.* 2016). Next, to identify genes critically responsible for *L. maculans* and *S. sclerotiorum* resistance in resistant accessions, we compared the *L. maculans* and *S. sclerotiorum* responsive chitinase genes in both resistant and susceptible accessions, respectively. Furthermore, we compared the 18 up-regulated chitinase genes on pathogen *L. maculans* aggression with 18 up-regulated chitinase genes against *S. sclerotiorum* attack. Interestingly, the upregulation after infection with *L. maculans* was stronger in resistant accession than in susceptible accessions. In contrast, the upregulation after *S. sclerotiorum* attack showed higher levels in both resistant and susceptible accessions whereas there was much less differences in in both resistant and susceptible accessions. In addition, there were some chitinase members that no expression changes was detected, which may be accounted for that some chitinases showed little or no enhancement of resistance and do not have any antifungal activity (WANG *et al.* 1999; TAIRA *et al.* 2005). The above results indicate that some members of the chitinase gene family have developed as powerful basal defence against various pathogens attack and other individual members of the chitinase gene family have evolved different roles in response to different environmental stresses in *B. napus*.

In summary, our study provides a comprehensive analysis of the chitinase gene family in the rapeseed, including gene identification, sequence features, physical location, evolutionary relationship, and expression patterns of chitinase genes responding to *L. maculans* and *S. sclerotiorum* infection, which could facilitate further dissection of the function of the chitinase gene family in rapeseed.

## 4. Materials and Methods

### 4.1 Identification of chitinase genes in *B. napus*

The v4.1 genome sequences and annotations of *B. napus* were downloaded from the FTP site of the Brassica database (ftp://brassicadb.org/Brassica_napus/)(CHENG *et al.* 2011). To identify chitinase genes in *B. napus*, Glyco_hydro_18 (PF00704) and Glyco_hydro_19 (PF00182) domains were obtained from the Pfam website (http://pfam.xfam.org/). The HMMER software version 3.0 was employed to identify chitinase genes against all known protein sequences (FINN *et al.* 2011). All candidate genes were further submitted to Pfam analysis (http://pfam.xfam.org/) to confirm the presence of one of the above two domains with E-value 0.0001. For annotation, the identified protein sequences were aligned with NCBI nr database using BLAST alignment (E-value cut-off of 1e-5) (ALTSCHUL *et al.* 1997). The identification of signal peptide was performed in the website (http://www.cbs.dtu.dk/services/SignalP/) (PETERSEN *et al.* 2011).

### 4.2 Phylogenetic tree construction, and sequence analysis

All identified chitinase genes were aligned using the MUSCLE program within MEGA 7.0 software (KUMAR *et al.* 2016). Subsequently, a neighbor-joining (NJ) method was then applied to construct a phylogeny of chitinase genes with a 1000 bootstrap replication. Motifs of chitinase proteins in *B. napus* were investigated statistically using online MEME software (http://meme-suite.org/tools/meme), which set the maximum number of motifs at 10. Subsequently, InterProScan (http://www.ebi.ac.uk/interpro/search/sequence-search) was employed to annotate the all identified motifs. In addition, the exon-intron structures of genes were performed with the gene structure display server program (http://gsds.cbi.pku.edu.cn/).

### 4.3 Chromosomal distribution and evolution patterns of chitinase genes

The chromosomal locations of chitinase genes were determined based on annotation data obtained from the *B. napus* database. The orthologous relationships between the chitinase genes in *B. napus* and *A. thaliana, B. rapa*, and *B. oleracea* genes were evaluated following the criteria: we used program BLAST to identify putative orthologues between chitinase genes in *B. napus* and one of *A. thaliana, B. rapa*, and *B. oleracea* species with both coverage over 70% and identity more than 70%, all chitinase sequences from *B. napus* was searched against all gene sequences from one of *A. thaliana, B. rapa*, and *B. oleracea* species. Tandem duplication was characterized as multiple genes of one family located within the same or neighboring intergenic region (LI *et al.* 2014).

### 4.4 Analysis of transcriptome sequencing data

The sequence data responsive to *L. maculans* infection was deposited in the BioProject Database of the National Center for Biotechnology Information under accession number PRJNA378851. Transcriptome data under accession number PRJNA274853 publicly available on the NCBI SRA database were mined and analyzed for expression patterns of the rapeseed chitinase genes in response to *S. sclerotiorum* infection. Sequencing reads were then aligned to the *B. napus* reference genome sequence (ftp://brassicadb.org/Brassica_napus/) using TopHat, v2.1.1 (KIM *et al.* 2013). Mapping data was used to estimate expression values for annotated genes using htseq-count tool (ANDERS *et al.* 2015). Differential gene expression analyses were performed using the R/Bioconductor package, DESeq2 (LOVE *et al.* 2014). An absolute value of log2 fold change >1.5 and the False Discovery Rate (FDR)< 0.05 was set to declare differentially expressed genes.

## Supplementary Materials

**Supplementary Figure 1.**
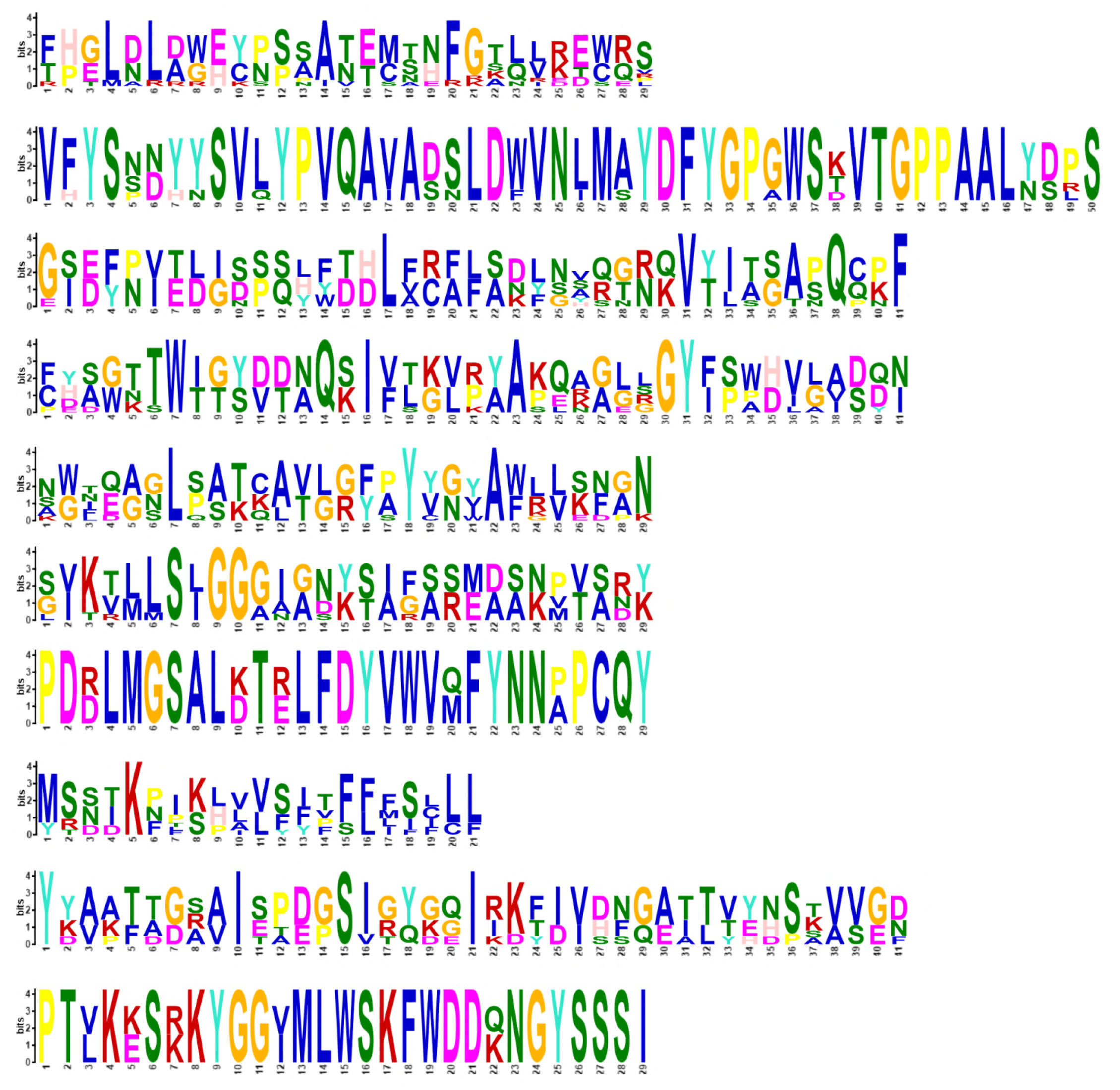
Details of the ten conserved motifs of chitinase GH-18 family as derived by MEME analysis.

**Supplementary Figure 2.**
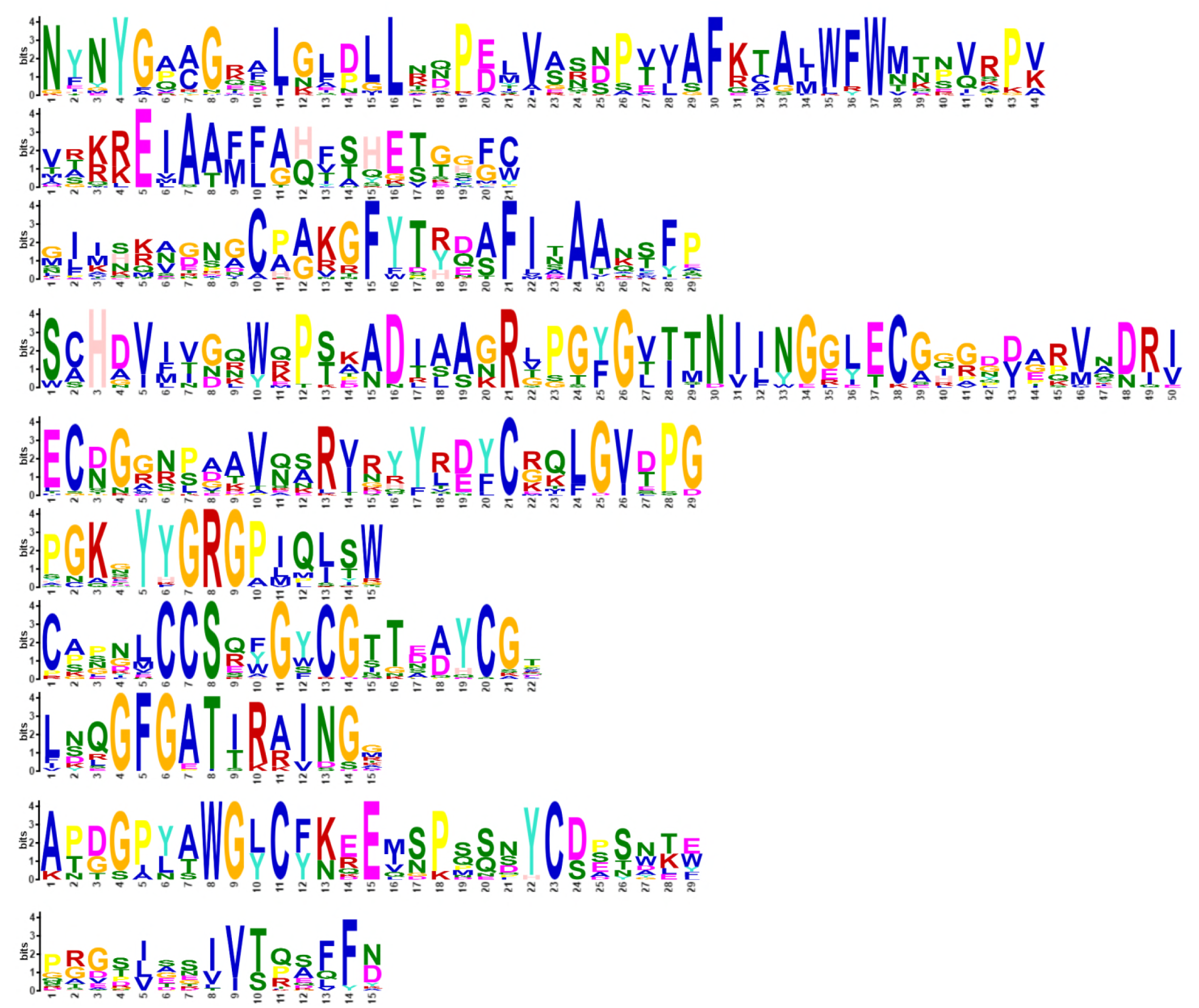
Details of the ten conserved motifs of chitinase GH-19 family as derived by MEME analysis.

**Table S1.**
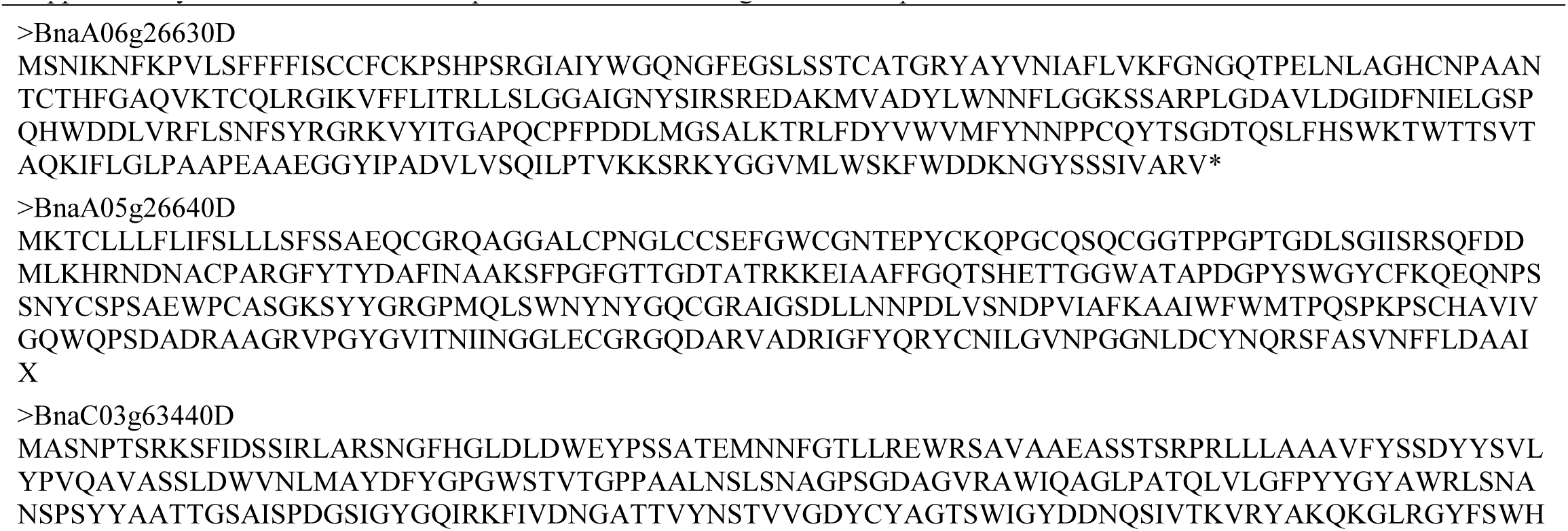
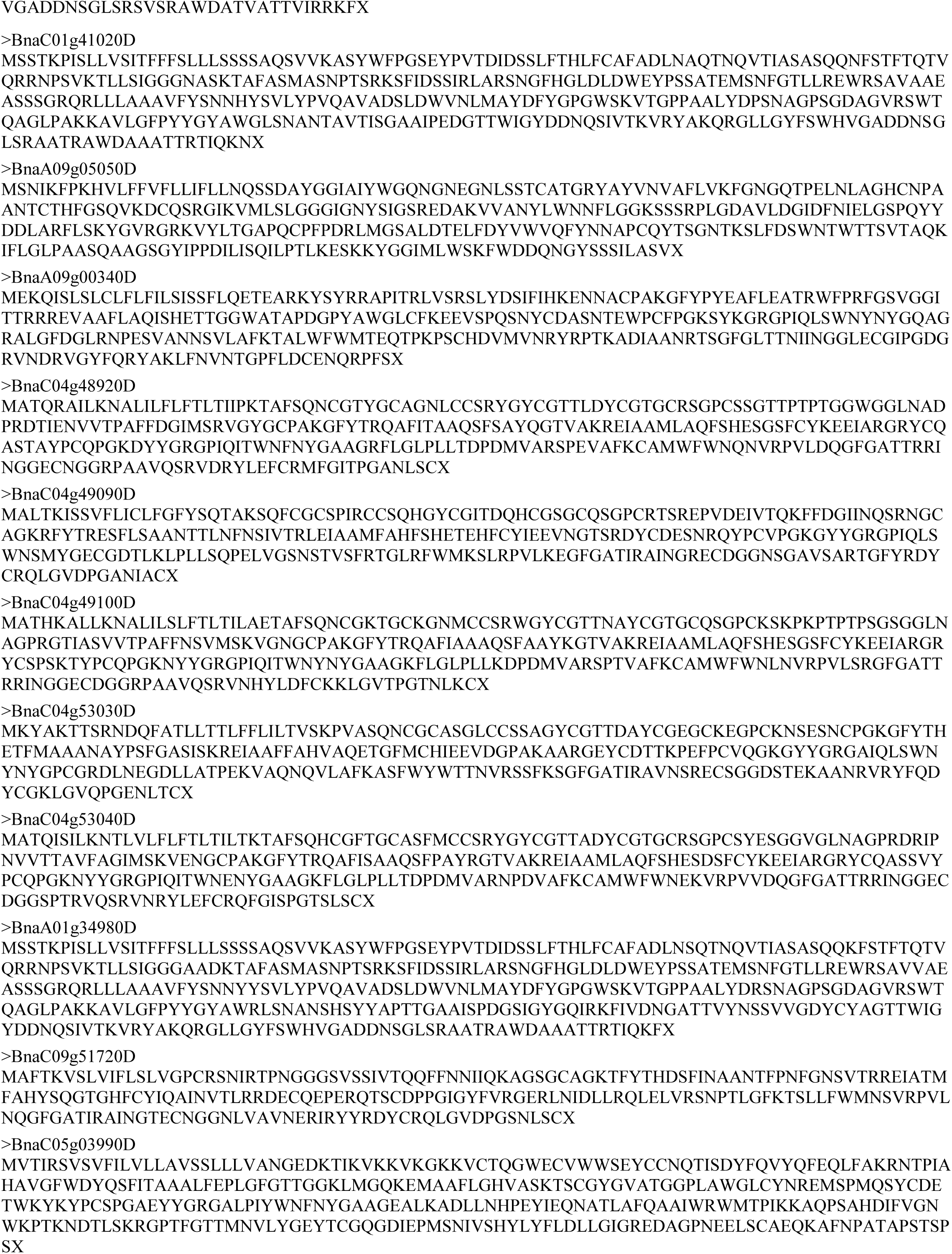
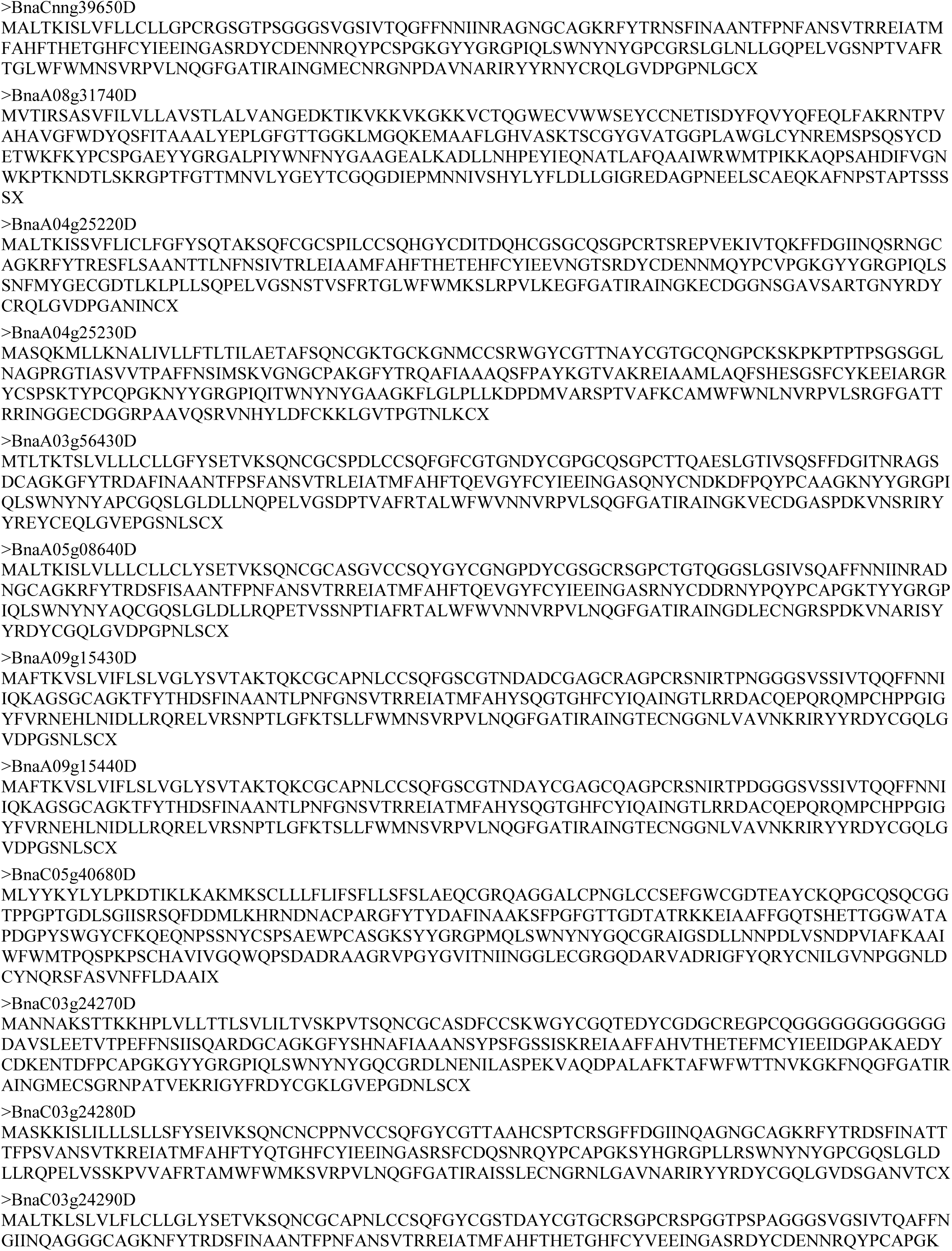
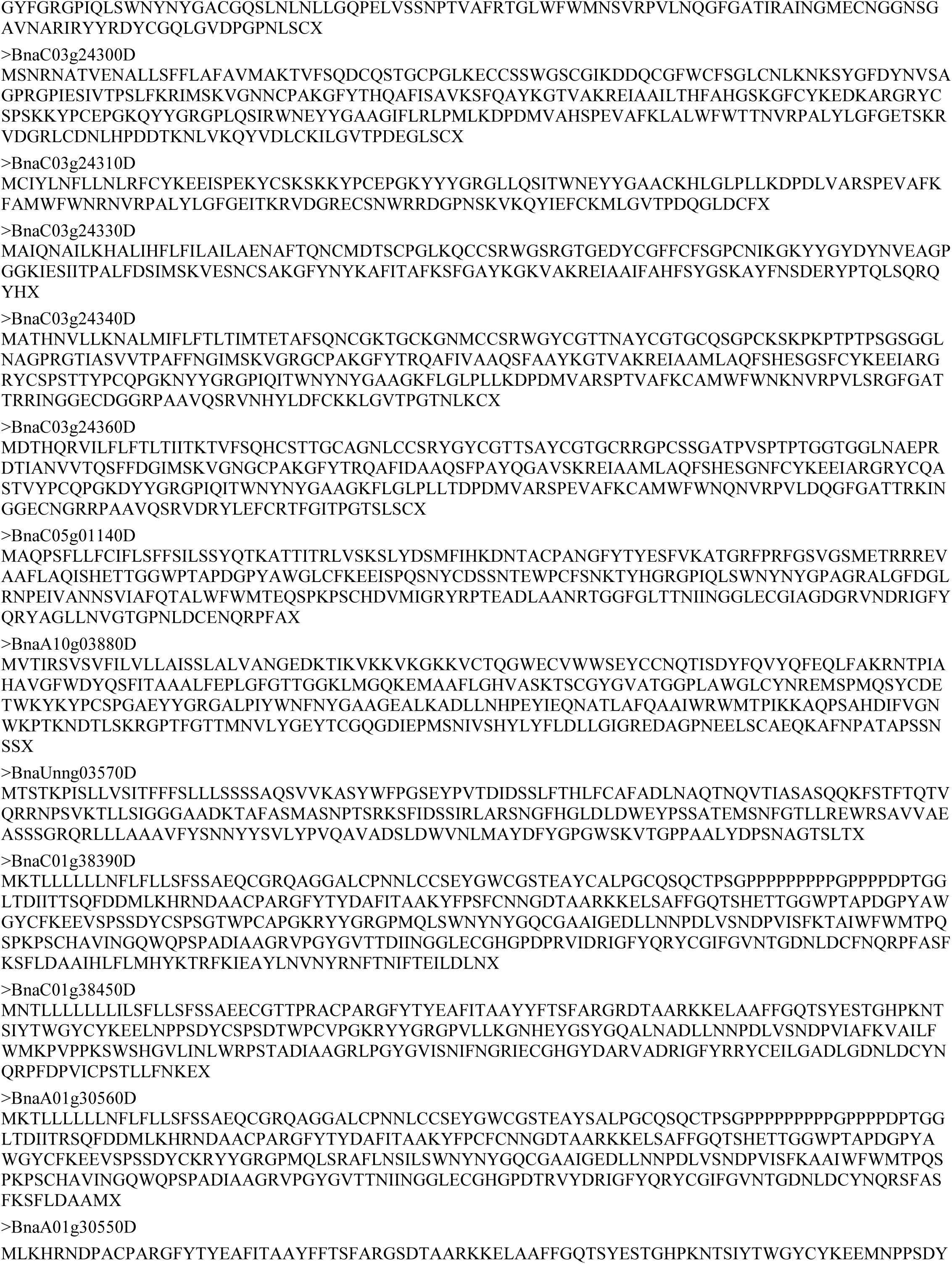
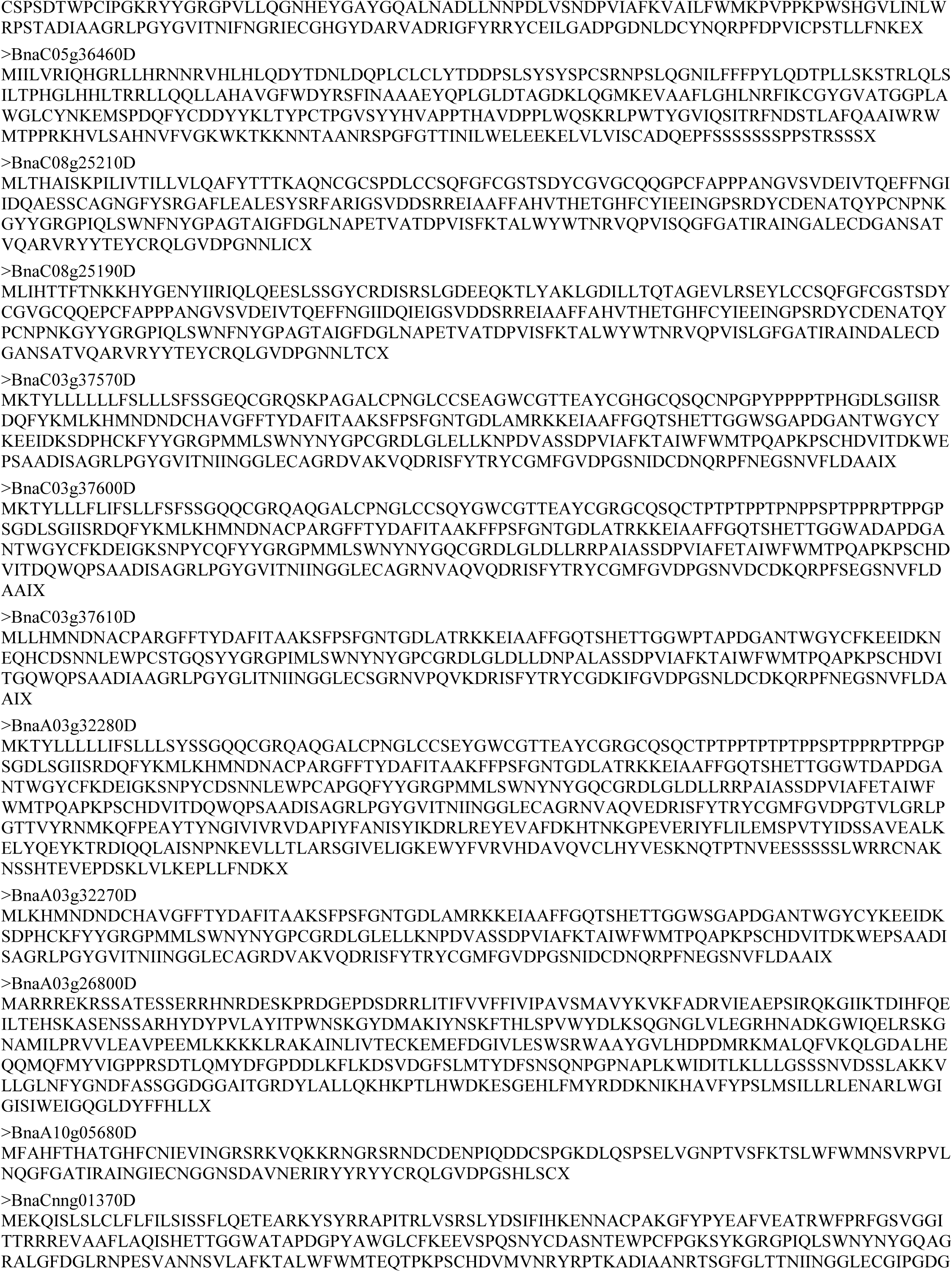
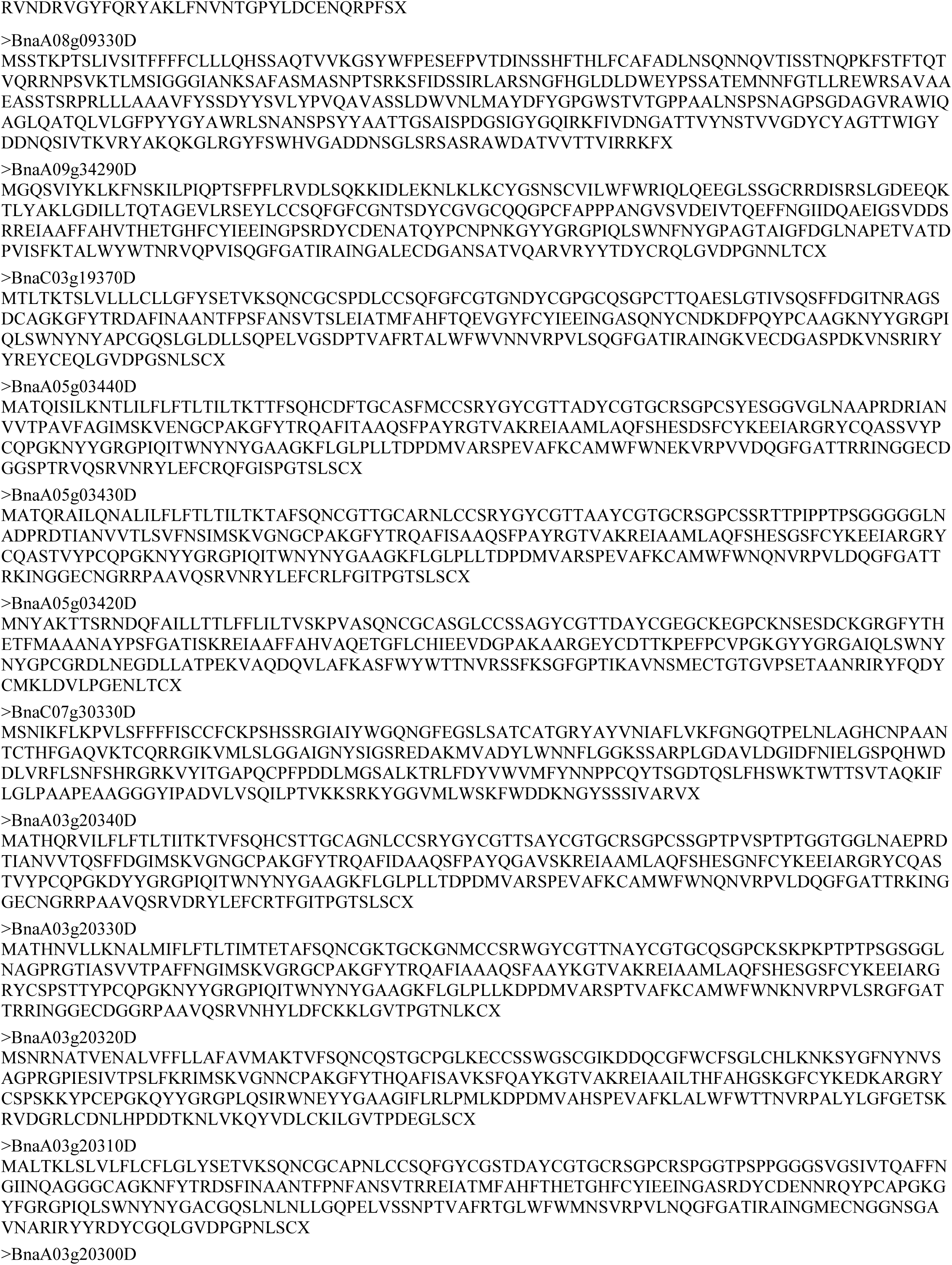
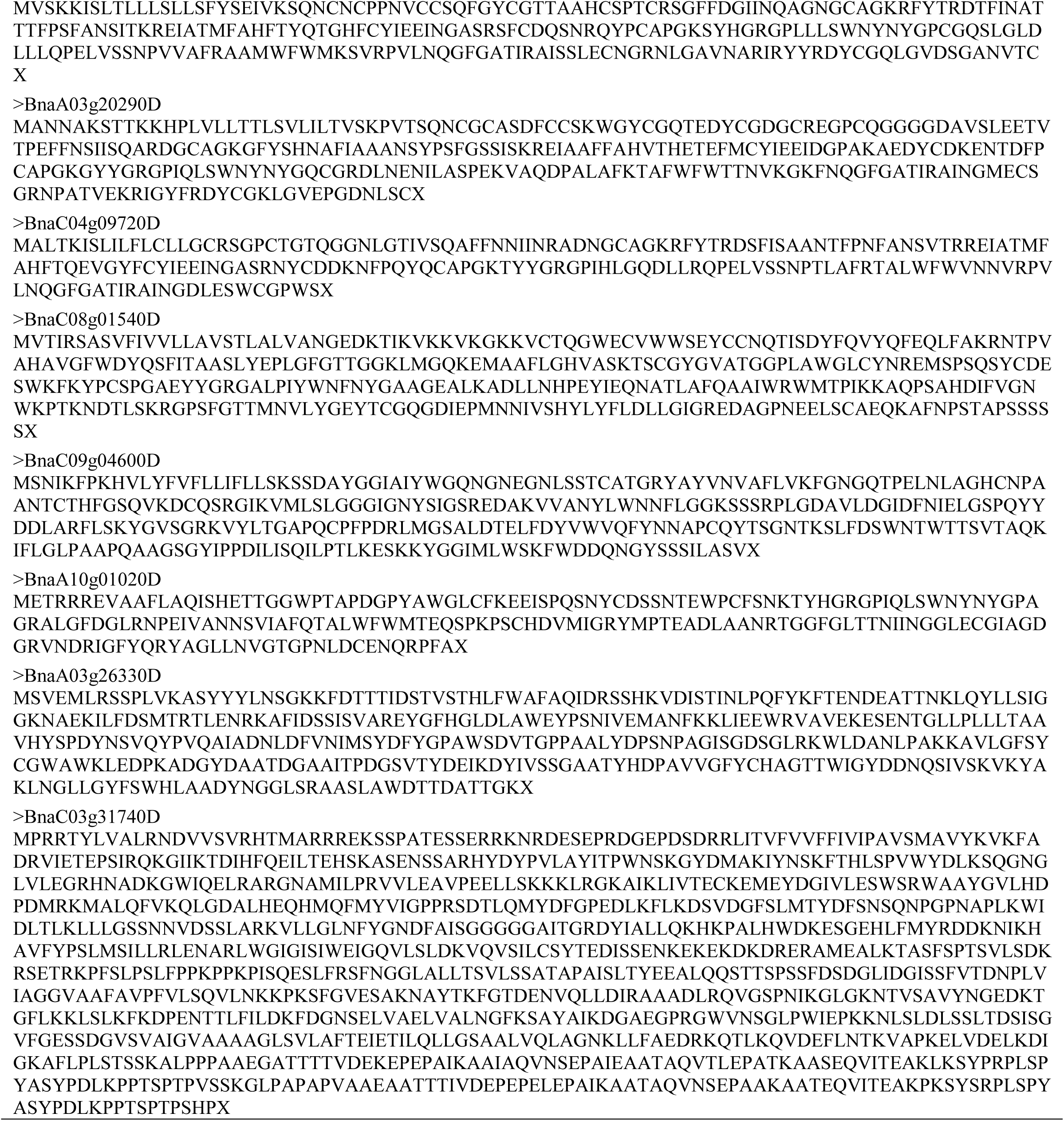
Amino acid sequences of 68 chitinase genes in *B. napus*

## Acknowledgments

This study was financially supported by the Independent Innovation Special Fund of Henan Academy of Agricultural Sciences (2018ZC78), the Henan Fundamental and Frontier Research Fund (162300410153) and by the Natural Sciences and Engineering Research Council (NSERC) CRD project and the Growing Forward project of SaskCanola and Agriculture and Agri-Food Canada (AAFC).

## Author Contributions

Wen Xu and Genyi Li designed the study. Tengsheng Zhou and Bo An performed the experiments. Wen Xu and Baojiang Xu analyzed the data and drafted the manuscript. Genyi Li and Wen Xu finished the manuscript. All of the authors carefully checked and approved this version of the manuscript.

